# CD11c+ myeloid cells are the predominant CD4+CCR5+ immune population in the foreskin and are increased in men with HIV-associated penile anaerobes

**DOI:** 10.64898/2026.05.11.724468

**Authors:** Lane B Buchanan, Yazan Khan, Jorge R Vargas, Zhongtian Shao, Victoria Menya Biribawa, Henry Rogers Ssemunywa, Annemarie Namuniina, Brenda Okech, Aaron AR Tobian, Daniel E Park, Cindy M Liu, Rupert Kaul, Ronald M Galiwango, Jessica L Prodger

## Abstract

Specific anaerobic species within the penile microbiome – Bacteria Associated with Seroconversion, Inflammation and Immune Cells (BASIC) – have been linked to increased HIV-1 susceptibility. These bacteria can directly disrupt epithelial integrity and are believed to increase local inflammation, resulting in an increased density of HIV-susceptible T cells in the inner foreskin. It is currently unknown whether other immune cells bearing the HIV entry receptors, CD4 and CCR5, are also elevated in individuals with a high abundance of BASIC species. Using inner foreskin tissues and penile swabs from males undergoing voluntary medical male circumcision, we performed a retrospective cross-sectional study to assess the relationship between BASIC species and the tissue density of such immune cells, including CD68+ macrophages, CD11c+ dendritic cells, and CD207+ Langerhans cells.

The most abundant cells in the inner foreskin expressing the HIV co-receptors were CD11c+ dendritic cells (48.6% of CD4+/CCR5+ cells), followed by CD68+ macrophages (28.6%), CD3+ T cells (18.8%), and CD207+ Langerhans-like (8.8%) cells. The absolute abundance of BASIC species was associated with elevated tissue densities of both CD4+/CCR5+ T cells (as previously reported) and a heterogeneous population of CD3-/CD4+/CCR5+ cells of myeloid origin. In the dermis, BASIC species abundance was linked to elevated densities of cells expressing CD11c, CD68, and CD207, as well as those co-expressing CD11c and CD207; furthermore, CD11c+ and CD207+ cells were farther from the basement membrane in participants with a high abundance of BASIC species. Myeloid cells were not elevated in participants with a high abundance of control taxa. In an integrated analysis including previously published data from this same cohort, myeloid-cell densities clustered tightly together, positively correlated with BASIC species and pro-inflammatory cytokines, and had trends to negative correlations with control taxa (significant for CD207+ cell density).

Overall, our findings suggest that BASIC species are associated with a broader foreskin immune phenotype marked by increased densities of HIV-susceptible myeloid and T cells, alongside epithelial disruption.

## Introduction

Voluntary medical male circumcision (MMC) decreases heterosexual HIV acquisition in men^1–3^, due at least in part to the reduction of penile anaerobic bacteria^4,5^. The abundances of six anaerobic species in the coronal sulcus microbiome – *Prevotella bivia*, *P. disiens*, *Peptostreptococcus anaerobius*, *Dialister micraerophilus*, *D. propionicifaciens*, and a genetic near-neighbour of *D. succinatiphilus* – have been linked to an elevated density of T cells bearing the HIV co-receptors CD4 and CCR5 in the inner foreskin, higher levels of pro-inflammatory cytokines, and the disruption of epithelial junction proteins in penile tissues^6–8^. These species are collectively referred to as Bacteria Associated with Seroconversion and Immune Cells (BASIC)^6^.

In the coronal sulcus and inner foreskin tissues BASIC species are thought to increase HIV acquisition through a combination of direct cleavage of epithelial junction proteins^8^ and by inducing local inflammation that leads to recruitment of HIV target cells to the inner foreskin, where they can become infected with HIV during sex^6^. In macaques, an elevated density of CD4+/CCR5+ T cells in the gut mucosa is a key determinant of whether exposure to simian immunodeficiency virus (SIV) results in infection^9^. In humans, elevated levels of genital inflammatory cytokines and chemokines are associated with increased risk of sexual HIV acquisition in both the penile coronal sulcus and the female genital tract^10–15^. Together, these data support a model in which microbial-driven mucosal inflammation increases HIV susceptibility by simultaneously compromising epithelial barrier function and increasing the local availability of HIV-susceptible cells.

A wide range of immune cells in penile skin co-express CD4 and CCR5, making them potential targets for initial HIV infection. Cells bearing HIV entry receptors – including T cells, Langerhans cells, dermal dendritic cells, and macrophages – are present throughout penile epithelia, with particularly high densities in the inner foreskin^6,16–21^. Langerhans cells are most superficial in the inner foreskin, placing them close to the epithelial surface and genital secretions, whereas macrophages and other dendritic cell subsets are more abundant in the underlying dermis^17,20,21^. In the vagina and rectum, *ex vivo* and *in vivo* models show that dendritic cells and macrophages, together with CD4+/CCR5+ T cells, are among the earliest HIV/SIV target cells^16,22^, and that myeloid cells can retain intact virions and pass them to T cells without necessarily undergoing productive infection themselves^18,23,24^. Anogenital dendritic cell subsets – especially CD11c+ – have emerged as particularly efficient HIV target and transmitter cells^19^. These studies suggest that multiple myeloid subsets, including Langerhans cells, dermal dendritic cells, and macrophages, are likely to contribute alongside T cells to the establishment of infection in penile skin.

While HIV-associated penile anaerobes correlate positively with CD4+/CCR5+ T-cell density, it remains unclear whether this relationship extends to other HIV-susceptible immune cells that are prominent in penile tissues, and whether these microbial communities are also associated with altered spatial positioning of myeloid cells relative to the epithelial barrier. Addressing these questions will improve understanding of how the local microbiome shapes penile HIV susceptibility and may help identify cellular targets for prevention strategies. In this study, we quantified the density and spatial localization of HIV-susceptible cell subsets in inner foreskin tissues from HIV-negative adult Ugandan men undergoing voluntary MMC. We focused on dendritic cells, macrophages, and Langerhans-like cells alongside CD4+ T cells and assessed how HIV-associated anaerobes relate to their density and positioning within foreskin tissue.

## Methods

### Study Population

This is a retrospective cross-sectional study using banked biological samples collected from 116 HIV-negative males who underwent voluntary MMC in Uganda in 2019. Samples were collected as part of a randomized controlled trial (RCT) examining the effect of antimicrobial agents on penile microbiota, immunology, and HIV susceptibility^6,25^. Of the 116 participants, 25 were randomized to the no treatment arm and did not receive any antibiotic (22%), while the remaining 91 men were randomized to a treatment course of either topical metronidazole (n=23; 20%), topical clindamycin (n=22; 19%), topical hydrogen peroxide (n=23; 20%), or oral tinidazole (n=23; 20%) before MMC. Analyses of the effect of antimicrobial treatment on the RCT primary endpoint (*ex vivo* HIV entry into inner foreskin-derived CD4+ T cells), as well as pre-specified secondary endpoints (abundance of BASIC and control taxa, soluble immune mediators, and T cell density) have been reported previously^7^. Previously published, publicly available data on the abundance of BASIC and control taxa, soluble immune mediators, and T cell density was accessed for this study^7^.

### Ethics Statement

All participants provided written informed consent for the future use of banked samples for research purposes. Soluble immune mediator and microbiome data were generated as a part of the parent RCT by the University of Toronto and George Washington University, respectively (University of Toronto Research Ethics Board RIS #35254; George Washington University Institutional Review Board #00000169), and de-identified data were provided to Western University June 1 2022. Quantitative immunofluorescent microscopy, and integration of bacterial, cellular, and soluble immune mediator data, was performed at Western University with approval from the Western University Human Ethics Research Board (Protocol 111441). De-identified tissue blocks were shipped to Western University in May 2022 and analyzed between 1 May 2022 and 28 February 2026.

### Biological Sample Collection and Processing

Detailed methods for the collection and analysis of penile swabs have been previously described^7^. Briefly, two inner foreskin swabs were collected by a study clinician immediately prior to surgical cleaning and MMC. Swabs were placed in a transport buffer comprised of PBS and a protease inhibitor (cOmplete, EDTA-free, Roche). Swabs were transported to the lab on ice and the eluant was aliquoted and stored at -80°C prior to assessment of bacteria and soluble immune mediators. Following MMC, the inner foreskin (the portion of the foreskin closer to the coronal sulcus which lies against the glans on the non-erect penis) was tagged with a suture and the tissue was immediately placed into room temperature R10 medium (RPMI 1640 media, 10% heat-inactivated fetal bovine serum, 10 U/mL penicillin, 10 μg/mL streptomycin, 250 ng/mL amphotericin B, and 2 mM L-glutamine; all Gibco, Invitrogen) and transported to the lab. Visible blood vessels and clots were excised and the inner and outer foreskin aspects were separated. Inner and outer foreskin sections of 0.25 cm^2^ were excised and placed into optimal cutting temperature (OCT) compound before freezing and storing at -80°C.

### Immunofluorescence Staining

Staining was performed as previously described^26^. Frozen foreskin tissue sections (10 µm thickness) were fixed for 10 minutes in 3.7% formaldehyde in 0.1 M PIPES buffer, pH 6.8 (Avantor). Sections were blocked with 10% normal donkey serum (Avantor), 0.1% Triton X-100 (Sigma-Aldrich) and 0.1% Sodium Azide (Sigma-Aldrich) in PBS. Details on the antibodies used are listed in **Supplemental Table 1**. Tissue sections were incubated with primary antibodies at 37°C for 1 hour (or at 4°C overnight) and with secondary antibodies at 25°C for 30 minutes. Sections were washed with PBS between incubations. Fluoromount-G Mounting Medium with DAPI (4′,6-diamidino-2-phenylindole, ThermoFisher Scientific) was used as a counterstain to visualize cell nuclei.

### Quantitative Immunofluorescent Microscopy

Tiled images of whole foreskin tissue sections were imaged (200×) using a DM5500B fluorescence microscope (Leica Camera AG) and exported as TIF files. The excitation and emission filters are listed in **Supplemental Table 2**. All subsequent analyses were performed on full tissue sections (as opposed to fields of view). Manual tracings of the epidermis were segmented in Fiji (version 1.54f). A custom workflow was designed to acquire the following regions using the epidermal tracings: apical edge, basal edge, and a 100 µm region of dermis from the basal edge. Surface areas of the entire epidermis and the 100 µm region of dermis were measured for cell density quantification.

Cells expressing the surface markers CD3, CD4, and CCR5 were quantified using a previously published^27^ machine learning cell segmentation workflow which uses the ImageJ StarDist plugin v0.3.0^28^. This methodology was then adapted to detect new cell surface proteins CD68, CD11c, and CD207. Fifty new 600 µm x 600µm training images^27^ were randomly selected and cells which express CD68, CD11c, or CD207 were manually traced. For these cells, we replaced the StarDist model architecture with SplineDist^29^, which better accommodates irregularly shaped cells. To train the adapted model, we configured the computing environment on Ubuntu (v20.04.6 LTS). NVIDIA Driver (Release 535) was used to enable GPU-accelerated model training. Anaconda (v23.3.1), TensorFlow (v2.11), scikit-image (v0.17.6), OpenCV (v4.5.3), CUDA Toolkit (v11.8.0), and cuDNN (v8.8.0) were installed. Package management was handled via pip (v22.1.2) and Snakemake (v7.32.4). The complete environment, including all the scripts and dependencies, is available at the public repository listed below. After cell segmentation from StarDist and SplineDist workflows, a supplementary manual correction workflow was implemented with the CellCounter tool in Fiji. Cells incorrectly identified by the model were removed, while missed cells were added. Co-expression of multiple cell surface markers was quantified using a custom script which detects overlapping regions between two markers.

A cell distance algorithm to analyze the shortest distance between segmented cells and select tissue regions was adapted from a publicly accessible script created by Michael Cammer from NYU Grossman School of Medicine^30^. The median distances to both the apical and basal edges were measured for each cell in the epidermis, and the median distance to the basal edge of the epidermis was measured for cells within the 100 µm dermal region. All scripts used in this study are available at the following repository: https://github.com/prodgerlab/SplineDistQuantification.

### Bacterial Quantification

Detailed methods for quantifying penile bacterial species have been previously described^7^. Total DNA was extracted from coronal sulcus swab eluent using a combination of chemical and enzymatic lysis. Penile microbiota were characterized by 16S rRNA gene–based amplicon sequencing (V3V4) and by broad-range real-time PCR (V3V4). Species-level classification for select genera was performed using an in-house Bayesian classifier trained by a curated training set. Using the resultant sequencing and qPCR data, absolute abundance of each penile taxon was calculated as the product of total bacterial load and proportional abundance of the taxa.

### Soluble Immune Mediator Quantification

Detailed methods for quantifying soluble immune mediators in inner foreskin swabs have been previously described^7^. An electrochemiluminescent-based detection system (MesoScale Discovery) was used. Samples were run in duplicate, and any sample with a coefficient of variation >30%, or that exceeded the upper limit of quantification, were re-run. Concentrations below the threshold of detection were assigned as the lower limit of detection, derived from the mean from the standard curves of all plates run.

### Data Analyses

#### Participant Grouping

Analyses were pre-defined to compare participants based on the combined absolute abundance of BASIC species or control taxa. BASIC species included: *Peptostreptococcus anaerobius, Prevotella bivia, Prevotella disiens, Dialister propionicifaciens, Dialister micraerophilus,* and a genetic near neighbour of *Dialister succinatiphilus*. Control taxa included: (i) two taxa that increase in absolute abundance after MMC (*Staphylococcus* and *Corynebacterium*)^4^ and (ii) two Gram-negative anaerobic taxa that are not associated with HIV seroconversion (*Negativicoccus* and *Helococcus*)^6^. The analytic cohort contained three mutually exclusive groups: “High BASIC” (n=21), “High Control” (n=21), and “No BASIC” (n=26). The “High BASIC” group was defined as individuals in the top quartile of BASIC species absolute abundance (5.26 log_10_ 16S rRNA gene copies/swab and above), but not in the top quartile of control taxa absolute abundance. The “High Control” group was defined as individuals in the top quartile of control taxa absolute abundance (5.40 log_10_ 16S rRNA gene copies/swab and above), but not the top quartile of BASIC species absolute abundance. The “No BASIC” group was defined as individuals with no detectable BASIC species without being in the top quartile of control taxa absolute abundance. The study size was determined by the availability of banked inner foreskin tissue sections with matched penile microbiome data from the parent RCT. Of 116 participants with available parent-study data, 68 met criteria for inclusion in the primary categorical analyses: High BASIC (n=21), High Control (n=21), and No BASIC (n=26).

#### Statistical Comparisons

Differences in demographics and bacterial load between bacterial groups were assessed using χ² tests of independence or Fisher’s exact tests where appropriate, and continuous variables were compared using Kruskal-Wallis tests. Differences in antimicrobial treatment distribution between bacterial groups were assessed using a χ² test of independence, with post-hoc pairwise comparisons conducted using Fisher’s exact tests and Bonferroni correction for multiple comparisons.

Immune cell densities were square-root transformed prior to modeling to better satisfy normality assumptions for statistical analyses. Differences in immune cell densities between participant groups (High BASIC, High Control, No BASIC) were assessed using generalized linear models with a Gaussian distribution and identity link. Primary models included bacterial group as the predictor. Multivariable models additionally included RCT treatment assignment and a treatment-by-bacterial-group interaction term to assess whether associations were consistent after accounting for prior antimicrobial treatment exposure.

Median cell distances to the apical edge of the epidermis were compared between groups by analysis of covariance (ANCOVA), with group as a fixed factor and thickness of the nucleated epithelium as a covariate. Median dermal cell distances to the basement membrane were compared between groups using Kruskal-Wallis tests followed by Dunn’s post-hoc tests. Graphs comparing participant groups were generated using GraphPad Prism (v.9.5.1).

#### Hierarchical Clustering Analyses

To determine if patterns of correlation exist among the various immune parameters that have been quantified in this cohort, immunofluorescence data generated as a part of this study was integrated with previously published immune data from the same cohort. Absolute abundances of BASIC species and control taxa were summed at the per-sample level. These abundances were log_10_-transformed, and, along with immune and epithelial variables, were standardized to z-scores (mean-centered and scaled to unit variance) prior to association testing (R 4.4.1)^31^. Pairwise associations were estimated as Spearman rank correlations using the rstatix package v0.7.2 (cor_mat, cor_get_pval; two-sided tests, 95% confidence)^32^. Multiple comparisons were controlled for via the Benjamini–Hochberg false discovery rate using base R stats (p.adjust, method = "BH"). To aid visualization and reveal block structure, hierarchical clustering was applied to the correlation matrix (excluding bacterial abundances), converting correlations to distances as d = (1 − r) / 2 and clustering with base R stats (hclust). Correlation heatmaps were generated with ggplot2 v3.5.2^33^.

## Results

### Participant Demographics

To examine associations between penile bacteria relevant to HIV acquisition and HIV target cell density (CCR5+/CD4+), participants were grouped based on absolute abundance of BASIC species or control taxa (*Staphylococcus*, *Corynebacterium*, *Negativicoccus*, and *Helococcus*). High BASIC participants (n=21) were in the top quartile of BASIC species abundance, but not in the top quartile for control taxa. High Control participants (n=21) were in the top quartile for control taxa, but not for BASIC species. This approach provided two groups of participants with similar overall bacterial load (median log_10_ 16S RNA copies/swab and IQR: High BASIC 6.74, 6.37–7.24; High Control 6.39, 6.05–9.92; not significant), but significantly different abundance of BASIC species (median log_10_ 16S RNA copies/swab and IQR: High BASIC 5.82, 5.42–6.10; High Control 3.49, 0–4.28; p<0.0001). A third group, the No BASIC group, were participants with no detected BASIC species (n=26). No BASIC participants had low overall bacterial density (median log_10_ 16S RNA copies/swab and IQR: 5.17, 4.81–5.43; p<0.0001 compared to either group) and undetectable levels of BASIC species^8^.

Detailed participant demographics and the effect of previous antimicrobial treatments have been described previously^7,8^. Mean age of participants was 26 years, 52% were married or co-habiting, 46% had more than one sexual partner in the past 6 months, 38% had vaginal sex in the last week, and 88% of participants reported washing their penis daily. Demographic and behavioural characteristics were comparable between groups, with no statistically significant differences between the High Control, No BASIC, and High BASIC groups. Treatment distribution differed significantly between bacterial groups (χ² test, p=0.025), consistent with the known impact of antimicrobial therapy on penile microbiota composition^7^. High BASIC participants were more likely to have received no treatment (n=10/21, 48%), and less likely to have received topical clindamycin (n=0/21, 0%), compared with No BASIC participants (2/26, 8%; p < 0.001 and 12/26, 46%; p = 0.004, respectively). No significant differences in treatment distribution were observed for the High Control group.

### BASIC species are associated with elevated CD3+ and CD3- HIV target cells in the inner foreskin

We first confirmed previously reported associations between BASIC species and HIV target cells. Inner foreskin tissue sections (n=21, 21, and 26 for High BASIC, High Control, and No BASIC groups, respectively) were stained for an immunofluorescence panel including CD3, CD4, and CCR5. Images of full tissue sections were used to quantify HIV target cells co-expressing CD4 and CCR5. Consistent with previous reports, participants in the High BASIC group had a higher density of T cells bearing the HIV co-receptors (8.15 × 10⁻^3^ √cells/μm²) compared to participants in both the High Control (5.77 × 10⁻^3^ √cells/µm², p = 0.007) and No BASIC groups (5.73 × 10⁻^3^ √cells/µm², p = 0.029) (**Figure 1**). There were no significant differences in CD3+/CD4+/CCR5+ cell densities between the High Control and No BASIC groups. To verify that the associations between BASIC species and HIV target cell density were not due to prior exposure to an antimicrobial treatment – beyond that of the effect of the treatment on bacteria (e.g., a result of topical application of a cream, etc.) – we used a multivariate generalized linear model that included prior antimicrobial treatment assignment. In this model, the difference in CD3+/CD4+/CCR5+ cell density between the High BASIC and High Control groups remained statistically significant (adjusted p-value = 0.020), but the association with the No BASIC group did not meet the threshold for statistical significance (adj. p = 0.091). There were no significant associations between CD3+/CD4+/CCR5+ cell density and antibiotic treatment assignment, after controlling for bacterial group.

**Figure 1.**
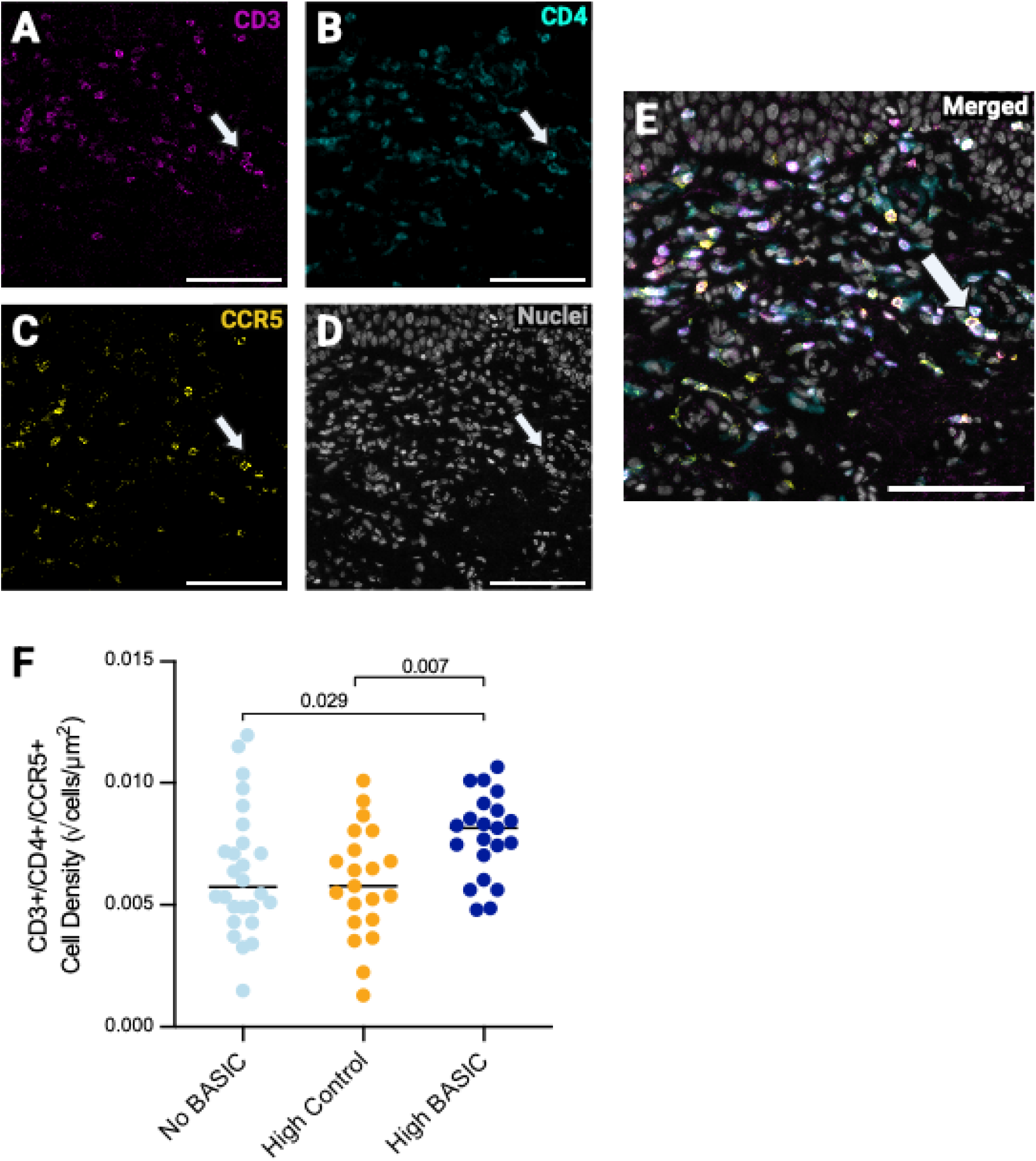
BASIC species are associated with elevated CD3+/CD4+/CCR5+ cells in inner foreskin tissue. Tissues were stained for the cell surface proteins CD3 (magenta, **A**), CD4 (cyan, **B**), and CCR5 (yellow, **C**), with nuclei counterstained in gray (**D**). Merged composite image is shown in **E**. Image scale bars are 100 µm. Brightness and contrast have been enhanced from the original images for visualization purposes. Cell densities were quantified from immunofluorescence images (**F**). Statistical comparisons were made between the High BASIC (n=21), High Control (n=21), and No BASIC (n=26) groups using a generalized linear model, α = 0.05. Lines represent the median in each group.

Participants in the High BASIC group also had an elevated density of CD3-/CD4+/CCR5+ cells (1.82 × 10⁻^2^ √cells/µm²) compared to both the High Control (1.11 × 10⁻^2^ √cells/µm², p = 0.0002) and the No BASIC groups (1.25 × 10⁻^2^ √cells/µm², p = 0.001) (**Figure 2**). No significant differences in CD3-/CD4+/CCR5+ cell densities were found between the High Control and No BASIC groups. The density of CD3-/CD4+/CCR5+ cells remained statistically significantly higher in men with High BASIC vs High Control (adj. p = 0.002) and No BASIC (adj. p = 0.019) after controlling for prior antibiotic treatment. There were no significant associations between prior treatment and CD3-/CD4+/CCR5+ cell density, after controlling for bacterial group.

**Figure 2.**
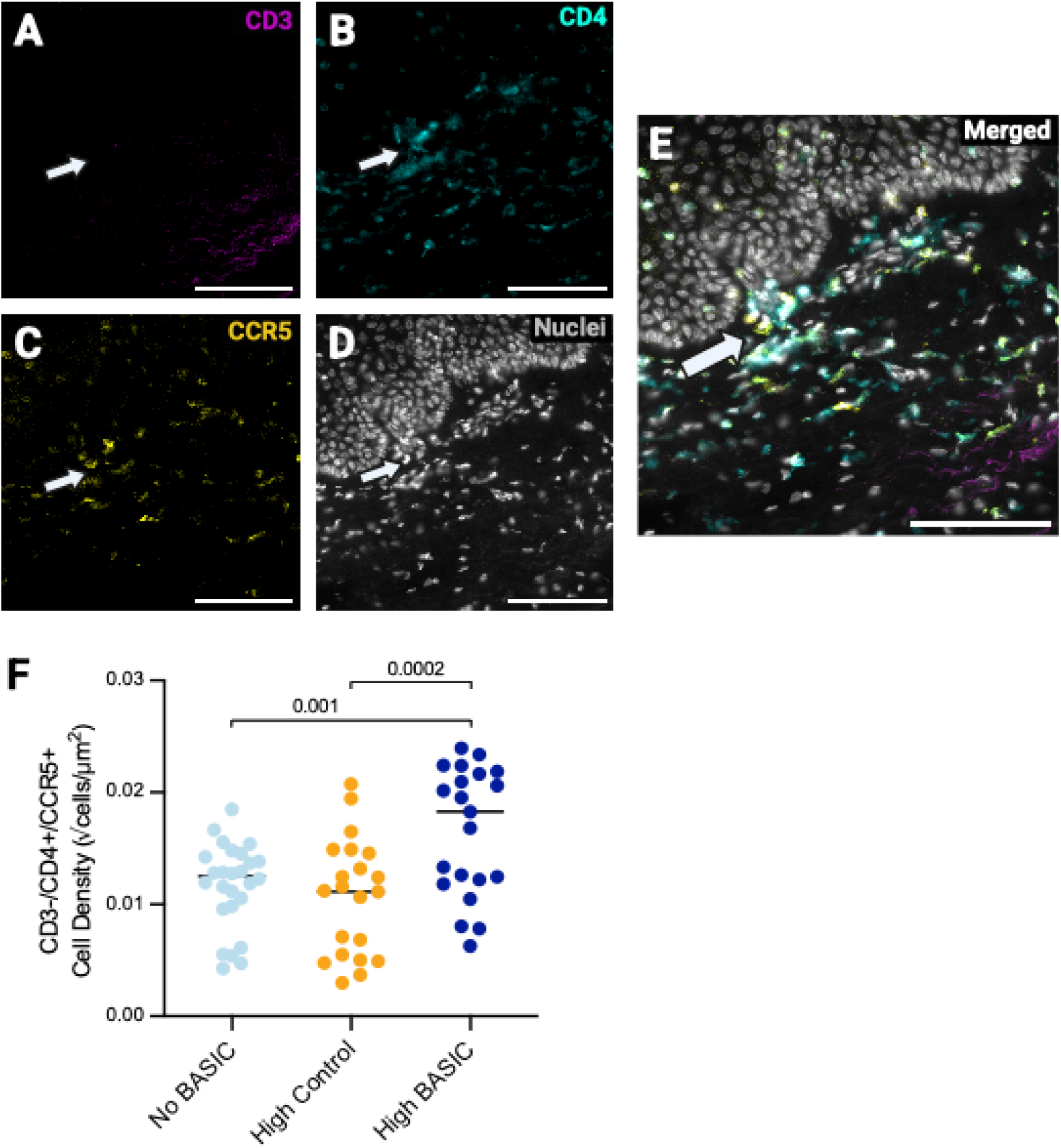
BASIC species are associated with elevated CD3-/CD4+/CCR5+ cells in inner foreskin tissue. Tissues were stained for the cell surface proteins CD3 (magenta, **A**), CD4 (cyan, **B**), and CCR5 (yellow, **C**), with nuclei counterstained in gray (**D**). Merged composite image is shown in **E**. Image scale bars are 100 µm. Brightness and contrast have been enhanced from the original images for visualization purposes. Cell densities were quantified from immunofluorescence images (**F**). Statistical comparisons were made between the High BASIC (n=21), High Control (n=21), and No BASIC (n=26) groups using a generalized linear model, α = 0.05. Lines represent the median in each group.

### Characterizing the association between BASIC species and CD3- HIV target cells

Several types of CD3- immune cells found in the foreskin^17,20,21^ have been reported to express the HIV co-receptors CD4 and CCR5 in both genital mucosa^19,34^ and human skin^35^. We assessed HIV co-receptor expression on putative dermal dendritic cells, macrophages, and Langerhans cells by staining a subset of tissues (5–6 tissues from different participants, 400x600 µm regions analyzed) through three additional immunofluorescence panels, each including the HIV co-receptors (CD4 and CCR5) alongside one cell surface marker: CD11c (**Figure 3A**), CD68 (**Figure 3B**), or CD207 (**Figure 3C**).

**Figure 3.**
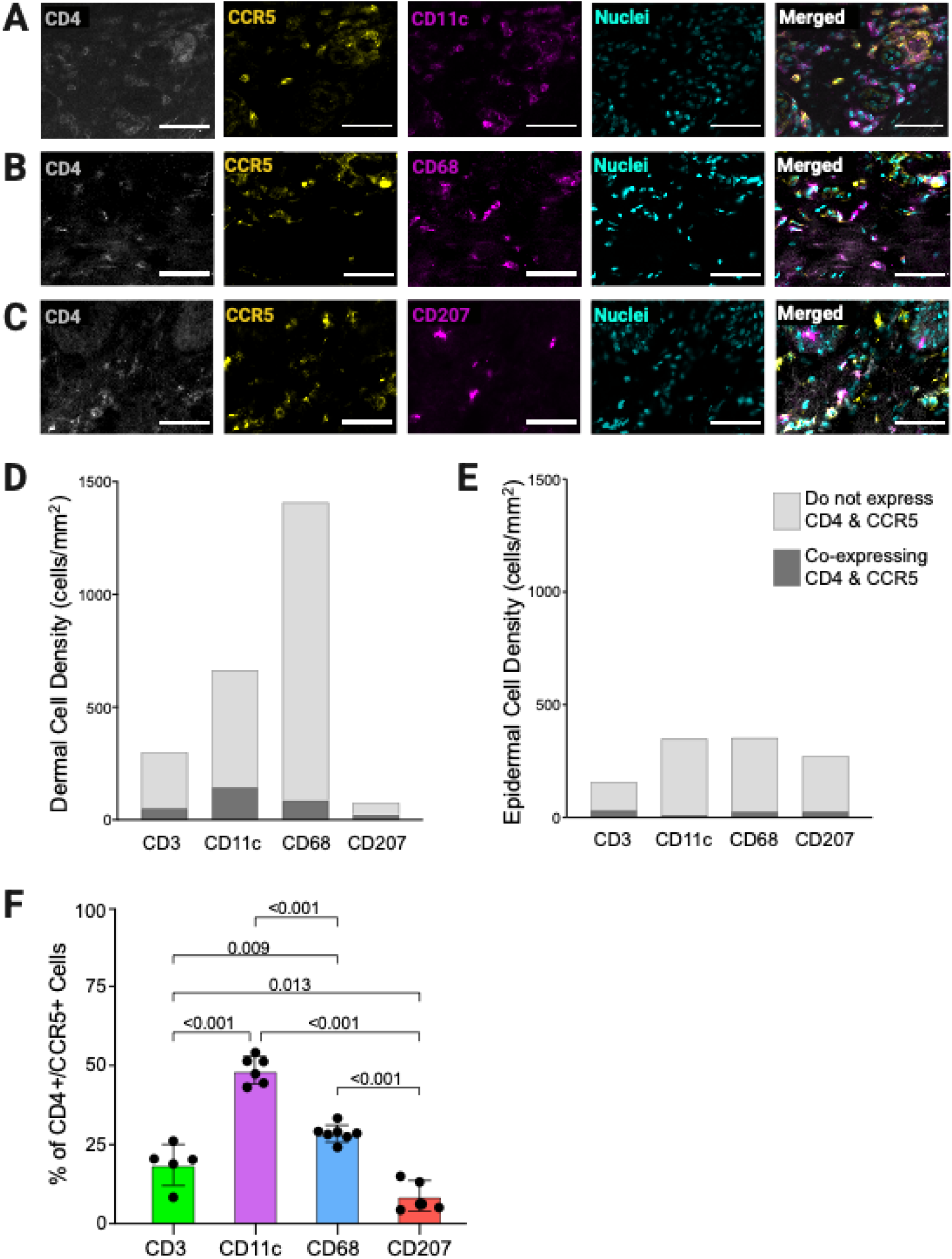
CD3- HIV target cells in inner foreskin tissue. Tissues were stained for the cell surface proteins CD4 (gray), CCR5 (yellow), and one of CD11c (**A**), CD68 (**B**), or CD207 (**C**) (magenta), with nuclei counterstained in cyan. Image scale bars are 50 µm. Brightness and contrast have been enhanced from the original images for visualization purposes. Mean cell density (cells/mm²) of each cell type was quantified in both the dermis (**D**) and epidermis (**E**), with the proportions of cells expressing HIV co-receptors overlaid onto each bar. The percent of CD4+/CCR5+ cells positive for each of the cell surface markers CD3, CD11c, CD68, and CD207 was quantified from immunofluorescence images (**F**). Lines represent the mean of n=5–6 FOV (from different participants) per cell type. Percentages of CD4+/CCR5+ cells were compared with ANOVA + Tukey’s test, α = 0.05.

In the dermis, CD68+ cells were the most abundant cell type (1409.6 cells/mm^2^), followed by CD11c+ (663.8 cells/mm^2^) and CD3+ (301.5 cells/mm^2^) cells (**Figure 3D**). CD207+ (78.0 cells/mm^2^) cells were comparatively rare in the dermis. In the epidermis, CD68+ (353.7 cells/mm^2^), CD11c+ (350.1 cells/mm^2^), and CD207+ (275.0 cells/mm^2^) cells were similarly abundant, with fewer CD3+ (158.6 cells/mm^2^) cells (**Figure 3E**).

However, the proportion of each cell type expressing the HIV co-receptors varied by cell type and location. Dermal CD207+ cells, while sparse, were most likely to co-express CD4 and CCR5 (32.0% of CD207+ cells), followed by CD11c+ (22.2%) and CD3+ (21.1%) cells. CD68+ cells were least likely to express HIV co-receptors (6.7%) (**Figure 3D**). In the epidermis, only a minority of CD11c+ and CD68+ cells expressed HIV co-receptors (2.1%, and 6.5%, respectively). In contrast, 23.9% of CD3+ cells and 19.3% CD207+ cells co-expressed CD4 and CCR5 (**Figure 3E**). Consequently, the most prevalent cell type in the inner foreskin expressing the HIV co-receptors was CD11c+ dendritic cells (48.6% of CD4+/CCR5+ cells), followed by CD68+ macrophages (28.6%), CD3+ T cells (18.8%), and CD207+ Langerhans-like (8.8%) cells (**Figure 3F**).

To determine which of these CD3- cell types is associated with BASIC species, inner foreskin tissue sections from all participants (n=21, 21, and 26 for High BASIC, High Control, and No BASIC groups, respectively) were stained for an immunofluorescence panel including CD11c, CD68, and CD207 (**Figure 4A**). Participants in the High BASIC group had a higher density of dermal CD11c+ cells (12.3 × 10⁻^3^ √cells/µm²) compared to the No BASIC (8.84 × 10⁻^3^ √cells/µm², p = 0.001) and High Control (9.43 × 10⁻^3^ √cells/µm², p = 0.042) groups (**Figure 4B**). These associations remained significant after controlling for prior treatment (vs No BASIC: adj. p = 0.005; vs High Control: adj. p = 0.013). Interestingly, we also observed that oral tinidazole treatment was associated with a significantly higher density of CD11c+ cells, after controlling for bacterial group (adj. p = 0.003).

**Figure 4.**
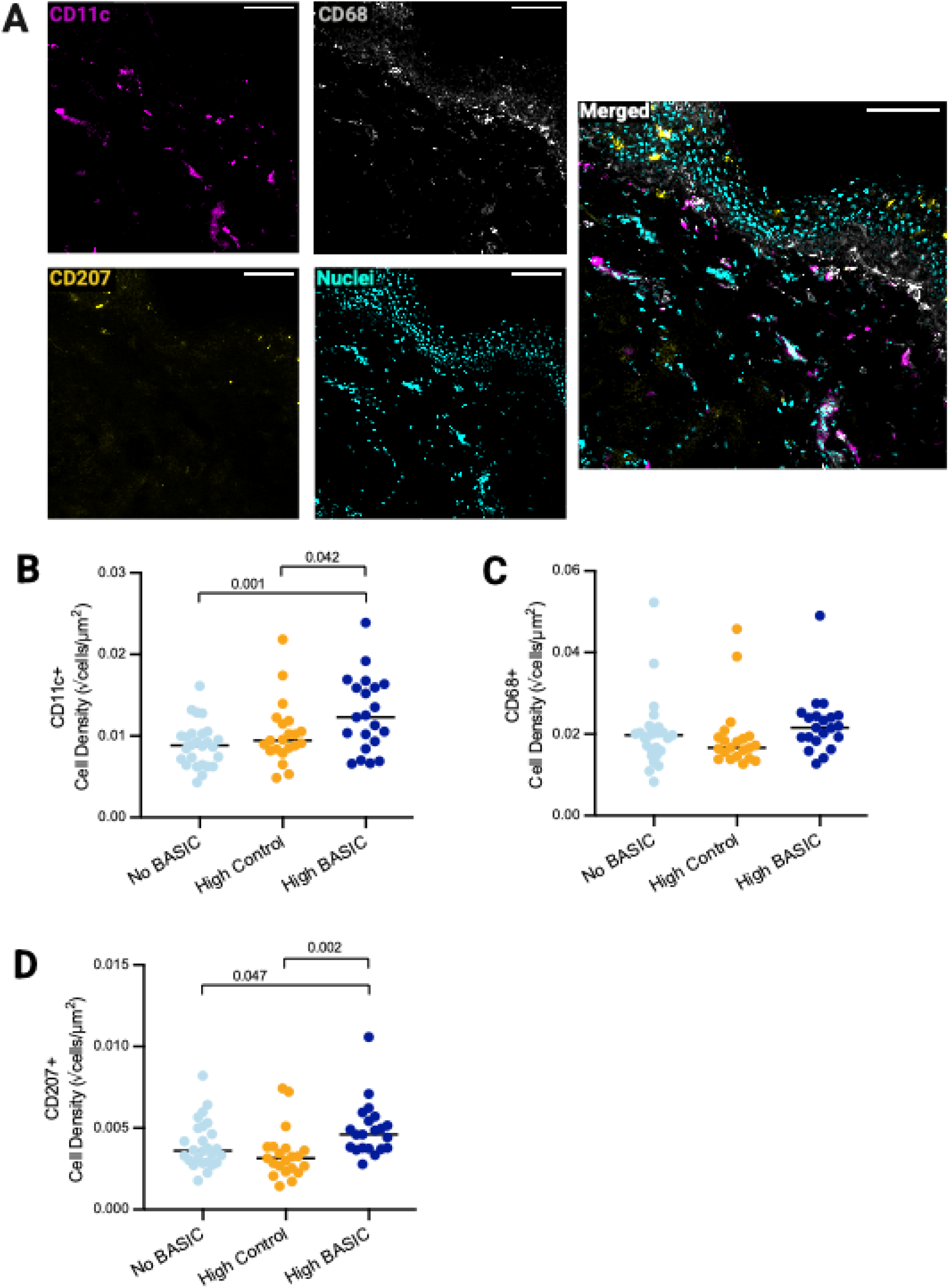
Density of CD11c+, CD68+, and CD207+ cells in the inner foreskin dermis. Tissues were stained for the cell surface markers CD11c (dendritic cells, magenta), CD68 (macrophages, gray), and CD207 (Langerhans cells, yellow), with nuclei counterstained in cyan (**A**). Image scale bars are 100 µm. Brightness and contrast have been enhanced from the original images for visualization purposes. CD11c (**B**), CD68 (**C**), and CD207 (**D**) cell densities were quantified from immunofluorescent images and statistical comparisons were made between the High BASIC (n=21), High Control (n=21), and No BASIC (n=26) groups using a generalized linear model, α = 0.05. Lines represent the median in each group. Merged composite image is shown in **E.**

In univariate analysis, there were no significant differences in dermal CD68+ cell density between the bacterial groups (**Figure 4C**). However, in the multivariate GLM including prior antimicrobial treatment, oral tinidazole was associated with significantly higher CD68+ cell density (adj. p = 0.002) and the High BASIC group had higher CD68+ cell density (21.6 × 10⁻^3^ √cells/µm²) compared to the High Control group (16.6 × 10⁻^3^ √cells/µm², adj. p = 0.044).

Finally, participants in the High BASIC group had significantly more dermal CD207+ cells (4.60 × 10⁻^3^ √cells/µm²) compared to the High Control (3.15 × 10⁻^3^ √cells/µm², p = 0.002) and No BASIC (3.61 × 10⁻^3^ √cells/µm², p = 0.047) groups (**Figure 4D**). The difference between High BASIC and Control groups remained significant after controlling for prior antimicrobial treatment (adj. p = 0.0017) and, like CD11c+ and CD68+ cell density, oral tinidazole was associated with significantly higher CD207+ cell density (adj. p = 0.010).

To better elucidate the identity of these myeloid cells, we compared cells co-expressing CD11c and either CD207 (**Figure 5A**) or CD68 (**Figure 5B**). Inner foreskin dermal CD11c+/CD207+ cells were elevated in the High BASIC group (2.20 × 10⁻^3^ √cells/µm²) compared to the High Control (1.43 × 10⁻^3^ √cells/µm², p = 0.004) and No BASIC (1.60 × 10⁻^3^ √cells/µm², p = 0.036) groups (**Figure 5C**). The difference between High Control and High BASIC groups remained significant after controlling for prior antimicrobial treatment (adj. p = 0.012). CD11c+/ CD68+ cells were also higher in the High BASIC group (7.60 × 10⁻^3^ √cells/µm²) compared to the No BASIC group (5.55 × 10⁻^3^ √cells/µm², p = 0.023). After adjusting for prior antimicrobial treatment, this association did not meet the threshold for statistical significance (adj. p = 0.066), but oral tinidazole treatment was associated with significantly higher CD11c+/CD68+ cell density (adj. p = 0.001). There was no significant difference in CD11c+/ CD68+ cell density between the High BASIC and High Control groups (**Figure 5D**).

**Figure 5.**
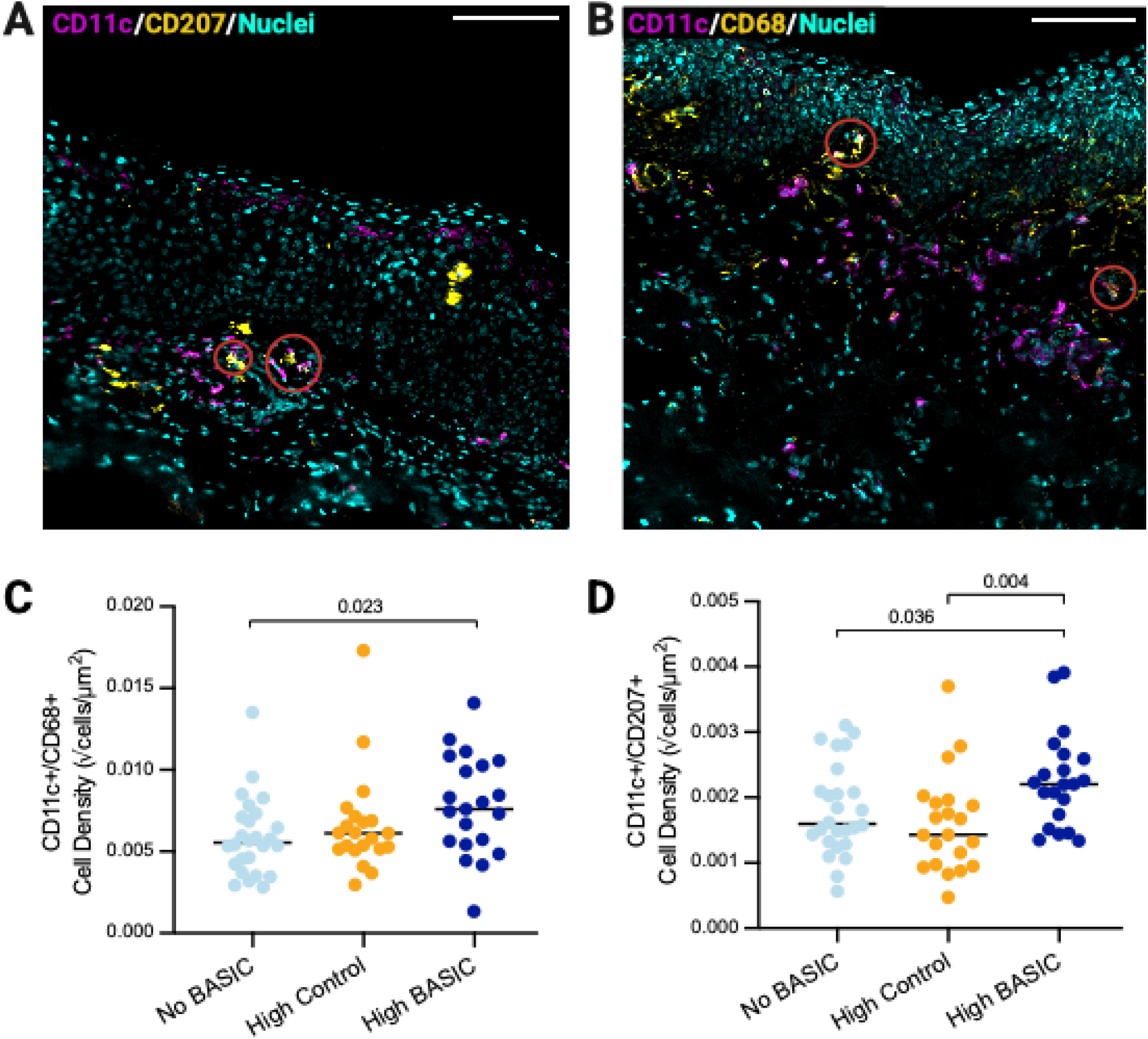
Density of CD11c+/CD68+ and CD11c+/CD207+ cells in the inner foreskin dermis. Tissues were stained for the cell surface markers CD11c (magenta) and CD207 (yellow) (**A**), or CD11c (magenta) and CD68 (yellow) (**B**), with nuclei counterstained in cyan. Image scale bars are 100 µm. Brightness and contrast have been enhanced from the original images for visualization purposes. CD11c+/CD207+ (**C**), and CD11c+/CD68+ (**D**) cell densities were quantified from immunofluorescence images and statistical comparisons were made between the High BASIC (n=21), High Control (n=21), and No BASIC (n=26) using a generalized linear model, α = 0.05. Lines represent the median in each group.

### No significant associations between BASIC species and epidermal density of CD11c+, CD68+, or CD207+ cells in the inner foreskin

In the epidermis, no significant differences were observed in CD11c+, CD68+, CD207+, or CD11c+/CD68+ cell densities between the High BASIC, High Control, and No BASIC groups (**Supplemental Figure 1**). Although not significant, the density of CD11c+/CD207+ cells was lower in the epidermis of the high BASIC group (5.36 × 10⁻⁶ vs 12.6 × 10⁻^6^ √cells/µm², p = 0.072) compared to the No BASIC group—an opposite trend to the association observed in the dermis.

### Epidermal CD68+ cells are closer to the apical surface in participants with a high abundance of control taxa

To determine the spatial localization of immune cells within the epidermis, an ImageJ algorithm was adapted from a publicly accessible script^30^ to measure the shortest distance from each cell to the apical or basal edge of the epithelium. Because BASIC species are known to correlate with increased epithelial thickness^8^, an analysis of covariance (ANCOVA) was performed with epithelial thickness included as a covariate to control for its potential confounding effect. After controlling for overall epithelial thickness, CD68+ cells were 8.81 µm closer to the apical edge of the epidermis in the High Control group compared to the No BASIC group (adj. p = 0.007). Mean cell distances to the apical and basal edges of the epithelium, as well as the model-adjusted group means after controlling for epithelial thickness, are shown in **Figure 6**.

**Figure 6.**
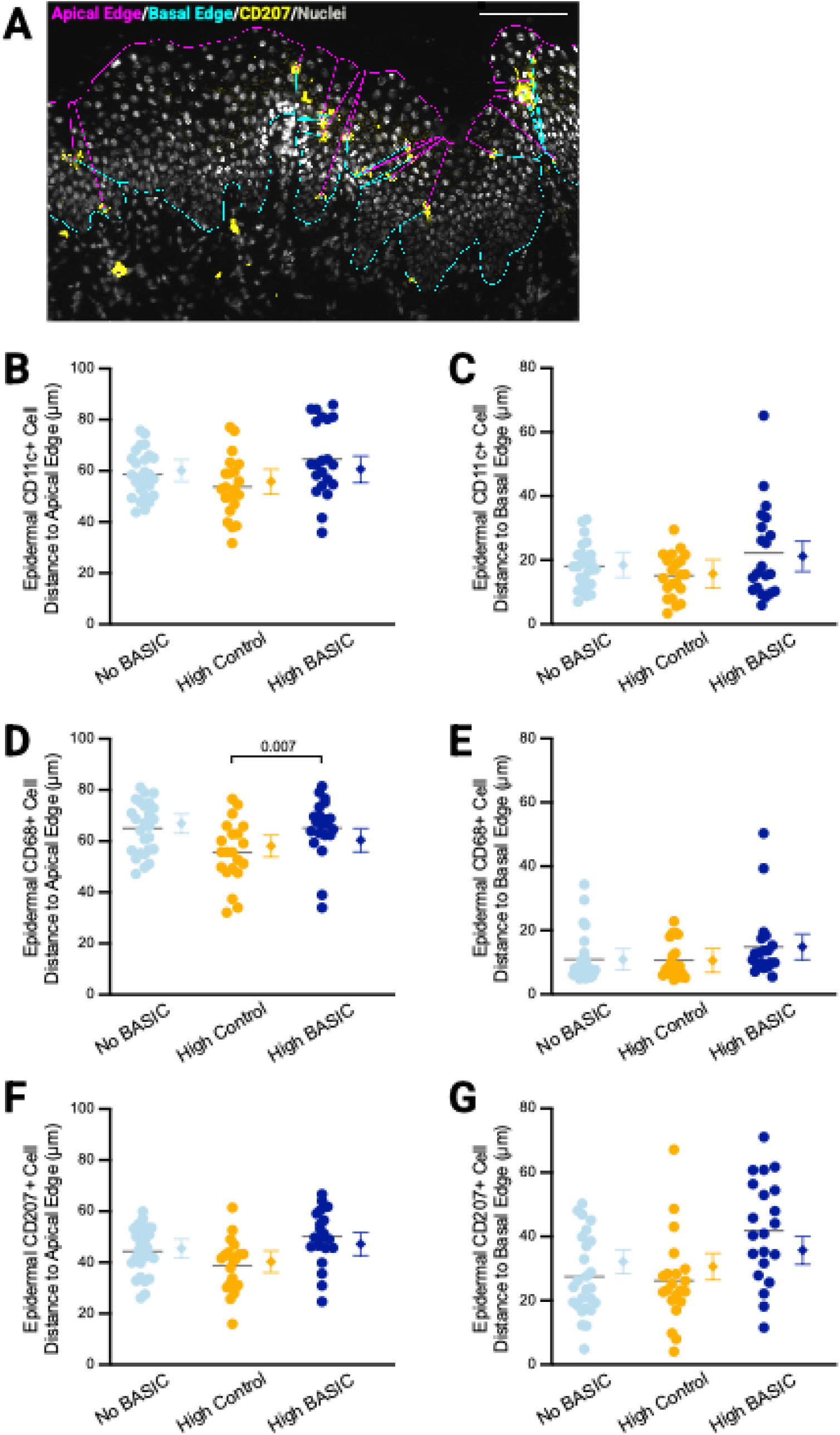
Distance of cells in the inner foreskin epidermis from the apical and basal edges. Cells were counted from immunofluorescence images and their distance to the apical (magenta) and basal (cyan) edges were quantified in ImageJ (**A**). Image scale bar = 100 µm. Distances to the apical and basal edges of the epithelium are shown for CD11c+ (**B**, **C**), CD68+ (**D, E**), and CD207+ (**F, G**) cells. Group means were compared between the High BASIC (n=21), High Control (n=21), and No BASIC (n=26) groups using analysis of covariance (ANCOVA) with epithelial thickness as a covariate, followed by pairwise comparisons of Tukey-adjusted estimated marginal means, α = 0.05. For each group, unadjusted distances for individual participant samples are shown with circles + median lines, while thickness-adjusted group means are shown with diamonds + 95% confidence intervals.

### Dermal CD11c+ and CD207+ cells are farther from the basement membrane in participants with a high abundance of BASIC species

The spatial localization of immune cells in the dermis was also assessed, by measuring the distance of each cell to the basement membrane (with an analysis area that included up to 100 µm into the dermis, measured from the basement membrane, **Figure 7A**). Dermal CD11c+ cells were significantly farther from the epidermis in the High BASIC group (median 31.6 µm) compared to the No BASIC group (26.7 µm, p = 0.026), with no significant difference observed compared to the High Control group (28.3 µm) (**Figure 7B**). Dermal CD207+ cells in the High BASIC group (40.9 µm) were significantly farther from the basement membrane compared to both the No BASIC (24.1 µm, p=0.0079) and High Control (23.3 µm, p=0.0065) groups (**Figure 7D**). No significant associations were observed between participant group and CD68+ cell distance from the basement membrane (**Figure 7C**).

**Figure 7.**
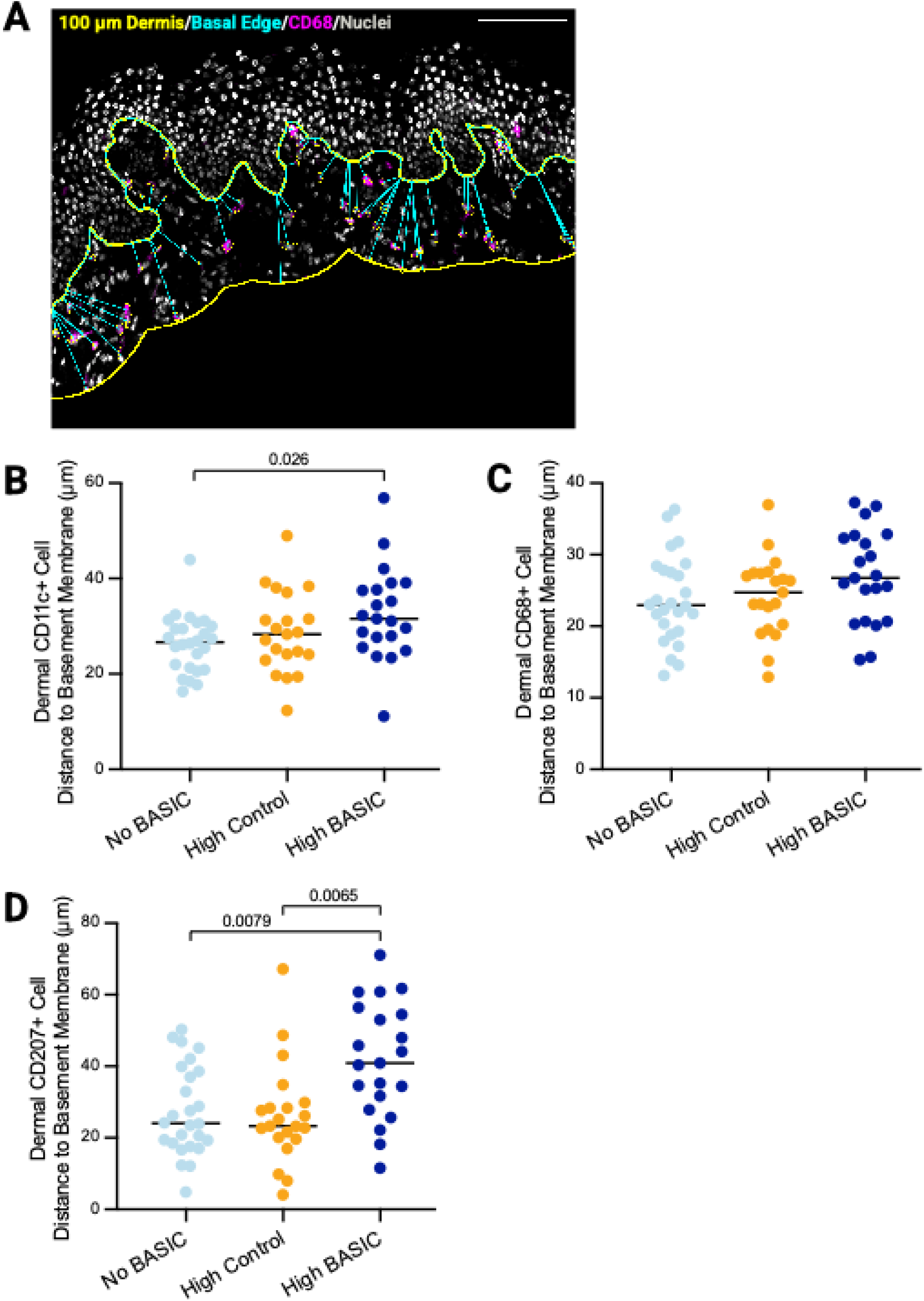
Distance of cells in the inner foreskin dermis to the epidermis. Cells were counted from immunofluorescence images and their distance to the basal edge of the epidermis (**A**) was quantified in ImageJ. Image scale bar = 100 µm. Distances are shown for CD11c+ (**B**), CD68+ (**C**), and CD207+ (**D**) cells. Statistical comparisons were made between the High BASIC (n=21), High Control (n=21), and No BASIC (n=26) groups using the Kruskal-Wallis test followed by Dunn’s post-hoc test, α = 0.05. Lines represent the median in each group.

### Antimicrobial treatments alter inner foreskin myeloid cell density

Because tissues were collected from participants enrolled in an RCT of antimicrobial treatments, we also examined associations between treatment assignment and immune-cell density, independent of bacterial grouping. Compared to men randomized to receive no treatment prior to circumcision, men who received oral tinidazole had significantly higher densities of dermal CD11c+ (11.56 vs 9.09 × 10⁻^3^ √cells/µm² p = 0.029), CD68+ (24.99 vs 17.51 × 10⁻^3^ √cells/µm², p = 0.002), and CD11c+/CD68+ (7.90 vs 5.69 × 10⁻^3^ √cells/µm², p = 0.006) cells, with a trend towards a higher density of CD11c+/CD207+ cells (2.06 vs 2.02 × 10⁻^3^ √cells/µm², p = 0.079). In contrast, men randomized to topical clindamycin or topical metronidazole had significantly lower densities of CD3-/CD4+/CCR5+ cells (12.10 × 10⁻^3^ √cells/µm², p = 0.030; 11.79 × 10⁻^3^ √cells/µm², p = 0.014, respectively) compared to men who received no treatment (18.17 × 10⁻^3^ √cells/µm²), and this effect was attenuated after accounting for bacterial group.

### Integrated analysis of the association between BASIC species and HIV target cell density, sub-preputial cytokines, and metrics of epithelial barrier integrity

The association between BASIC species and CD3+ T cell subsets, soluble immune mediators in the sub-preputial space, and markers of epithelial integrity have been previously reported, for this and other cohorts^6–8^. However, patterns in the expression of these various immune parameters have not been previously explored. We used hierarchical clustering and a correlation matrix to integrate all immune parameters that have been measured on participants of this RCT (n=116), including cell densities, soluble mediator concentrations, epithelial and *stratum corneum* thicknesses, and tissue expression of the epithelial junction proteins (E-cadherin, claudin-1, and desmoglein-1). This approach revealed six distinct clusters of immune markers (**Figure 8**). Broadly, two clusters were defined by epithelial metrics, two by immune cell types, and two by soluble immune mediators.

**Figure 8.**
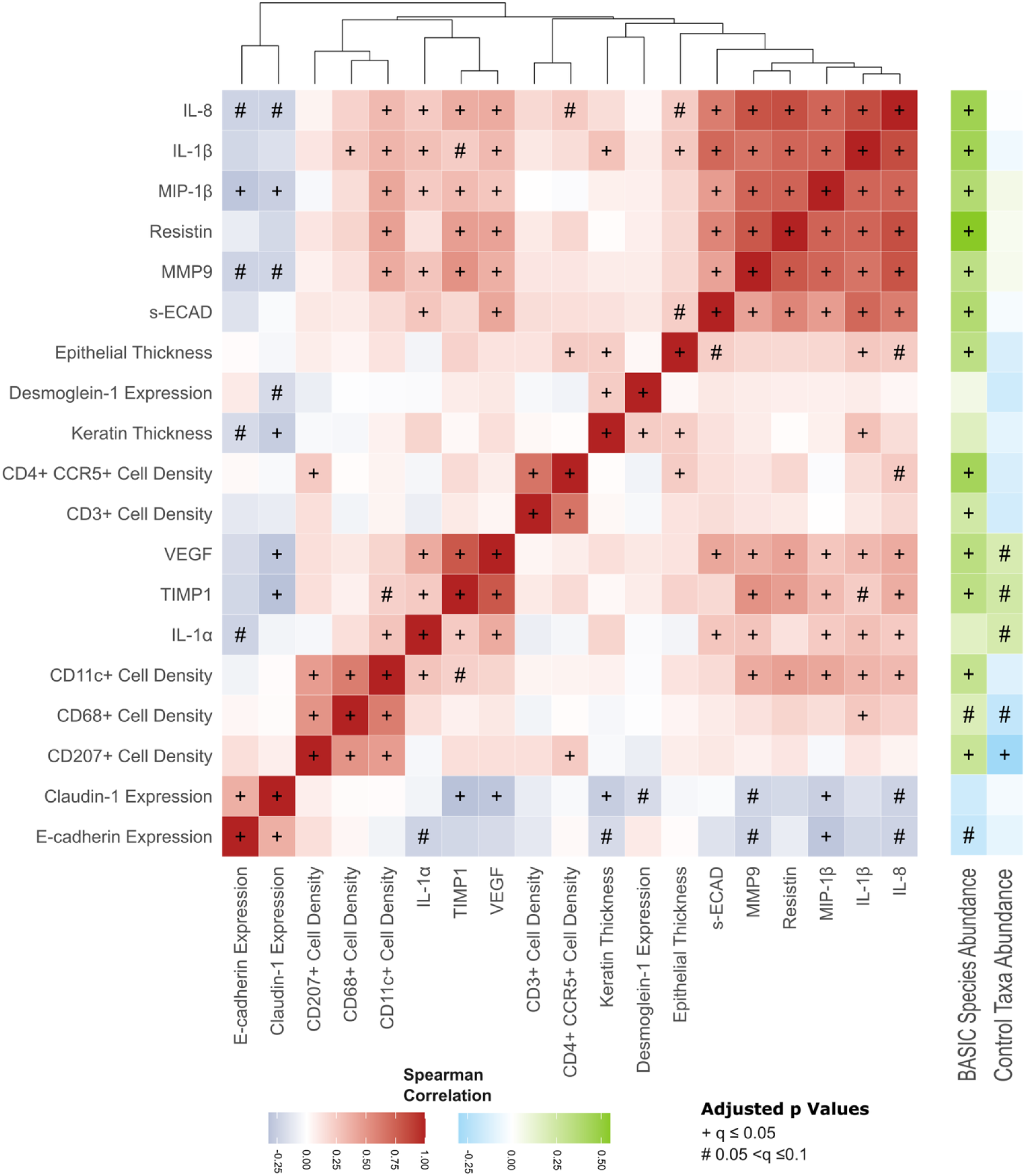
Hierarchical clustering of correlations between HIV target cells, soluble immune mediators, epithelial parameters. Soluble immune mediators were measured in sub-preputial penile swab eluents (n=116) by multiplexed immunoassay. Quantitative immunofluorescence was performed on inner foreskin tissues (n=116) to immune cell populations, epithelial thickness, keratin thickness, and epithelial junction protein expression. Hierarchical clusters are shown at the top. Correlations with abundances of BASIC species and control taxa are shown on the right; bacterial abundances were not included in the hierarchical clustering analysis. Significant Spearman’s correlations (α = 0.05) are denoted by +, trends (α = 0.1) are denoted by #.

Two contrasting profiles defined by epithelial metrics were observed: expression of E-cadherin and claudin-1 were tightly positively correlated, while desmoglein-1 expression was positively correlated with stratum corneum thickness (or “keratin” thickness; defined by filaggrin staining), with an inverse relationship between these two pairs of markers. Tissue E-cadherin expression was negatively correlated with BASIC species (no association with control taxa), and together E-cadherin and claudin-1 were additionally negatively correlated with most soluble immune mediators, IL-1α, TIMP1, VEGF, MMP9, MIP-1β and IL-8. In contrast, desmoglein-1 and stratum corneum thickness were not associated with BASIC species or control taxa, and had weakly positive (non-significant) associations with soluble immune mediators.

Two immune profiles (clusters) were defined by sets of positively correlated soluble immune mediators. One was defined by tight correlation of IL-1α, TIMP1, and VEGF, and the other by tight correlation of soluble E-cadherin (generated by cleavage of tissue E-cadherin), IL-8, IL-1β, MIP-1β, Resistin, and MMP9, and epithelial thickness. These two profiles were related, with weak positive correlations across the two profiles, and both were positively associated with the abundance of BASIC species. However, IL-1α, TIMP1, and VEGF also trended towards being positively correlated with the abundance of control taxa (p<0.01), while sE-cadherin, IL-8, IL-1β, MIP-1β, resistin, and MMP-9 were not.

Finally, two profiles were defined by correlated immune cell types: CD11c+, CD207+, and CD68+ cells were positively correlated with one another, while the overall abundance of CCR5+/CD4+ cells was more tightly correlated with CD3+ cell density. Both profiles were positively associated with BASIC species abundance. While the density of T cells was not significantly associated with immune mediator concentrations, the density of CD11c+ cells was significantly associated with increased concentrations of IL-1α, IL-1β, IL-8, MIP-1β, resistin, MMP-9 and soluble E-cadherin (all p<0.05). Interestingly, the only significant correlation with abundance of control taxa was a negative correlation with the density of CD207+ cells (p<0.05) and a trend towards a negative correlation with the density of CD68+ cells (p<0.01).

## Discussion

Our findings support a model in which HIV-associated penile anaerobes are linked not only to increased densities of CD4+/CCR5+ T cells, but also to a broader HIV-susceptible immune phenotype in the inner foreskin, encompassing both CD4+/CCR5+ T cells and multiple dermal myeloid populations. In particular, we found that myeloid cells accounted for most CD4+/CCR5+ cells in foreskin tissue, with CD11c+ cells representing the largest single fraction. We further found that control taxa were not positively associated with foreskin myeloid-cell density (tending toward negative associations), consistent with a more homeostatic tissue phenotype.

We have previously reported that CD4+/CCR5+ T cells are elevated in participants with a high abundance of HIV-associated penile anaerobes,^6^ and that reducing these anaerobes correspondingly reduces penile inflammation and CD3+ HIV target cells^7^. The current study extends this work by showing that BASIC species are associated not only with T-cell enrichment, but also with increased densities of CD3- HIV target cells. These cells were enriched for CD11c+ dendritic cells, with smaller contributions from CD68+ macrophages and CD207+ Langerhans-like cells, indicating that BASIC species are associated with a broader increase in HIV-susceptible myeloid populations rather than with T cells alone. By contrast, control taxa were not positively associated with T or myeloid target-cell density and, in some analyses, were negatively associated with macrophage and Langerhans-like cell density. Together, these findings suggest that BASIC species are associated with a distinct penile immune landscape characterized by epithelial barrier disruption^8^ and enrichment of multiple HIV-susceptible cell types.

Earlier histological studies of the foreskin identified abundant HIV target cells in the inner foreskin, including Langerhans cells, dermal dendritic cells, macrophages, and CD4+ T cells, with particularly high densities near the epithelial surface of the inner foreskin^17,20,21,36,37^. Together with ex vivo infection models and circumcision trials, these studies established the inner foreskin as an HIV-susceptible tissue, but the relative contributions of different myeloid and T-cell subsets to transmission remain debated^1–3,38–41^. Macaque SIV vaginal and rectal challenge models show that early infection is strongly biased toward CCR6+ CD4 T cells, with additional targeting of CD3-dendritic cells in the rectum, even though these subsets are relatively rare^16,22,42^.

Myeloid cells may also contribute to establishing infection by capturing virus and passing it to T cells, even when they are not themselves the predominant productively infected population. Bertram et al. identified a population of CD11c+ epidermal dendritic cells in human anogenital tissue that are more abundant than Langerhans cells, express higher levels of CCR5, preferentially interact with HIV, support higher levels of viral uptake and replication, and are more efficient at transmitting virus to T cells than Langerhans cells^19,43^. Buffa et al. further synthesize these and other data, highlighting mucosal dendritic-cell subsets – particularly CD11c+ dendritic cells – as key mediators of HIV capture and transfer to T cells in the anogenital tract^19,43^. Our findings build on this literature by showing that CD11c+ cells comprise the largest fraction of CD4+/CCR5+ cells in the inner foreskin, and that HIV-associated anaerobes are linked to increased densities of these cells.

The spatial distribution of myeloid cells was also altered. In participants with a high abundance of BASIC species, dermal CD11c+ and CD207+ cells were located farther from the foreskin basement membrane, suggesting that BASIC species are associated with remodelling of myeloid cell positioning within the foreskin. One possible explanation is that exposure to BASIC-associated inflammation alters myeloid-cell trafficking or maturation state, although these mechanisms were not directly measured here. Increased environmental sampling by inner foreskin myeloid cells has been linked to enhanced HIV transfer to dermal T cells in human foreskin explants^40,41,44^. More broadly, our findings indicate that HIV-associated anaerobes are linked not only to immune-cell density, but also to tissue organization.

Hierarchical clustering analysis suggested that densities of myeloid cells are tightly correlated with one another and with pro-inflammatory cytokines. More broadly, higher BASIC species abundance was associated with a composite phenotype characterized by disrupted epithelial barrier features, elevated soluble immune mediators, and increased densities of multiple HIV-susceptible cell subsets. This integrated pattern may help explain why differences in BASIC-species abundance are linked to differences in HIV risk^6^. By contrast, *Corynebacterium* and other control taxa were associated with lower myeloid-cell density (significant for CD207+ cell density) and preserved epithelial integrity (no correlation between their density and soluble E-cadherin).

Unexpectedly, oral tinidazole assignment was associated with higher densities of several dermal myeloid-cell populations. In the parent trial, oral tinidazole did not reduce abundance of coronal sulcus BASIC species, and controlling for bacterial group in the present study did not alter the positive association between tinidazole treatment and myeloid cell density. These findings raise the possibility that oral tinidazole has effects on foreskin myeloid-cell density that are not fully explained by the measured bacterial groupings used in this study.

Based on these findings and previous publications, we propose a mechanistic model in which HIV-associated penile anaerobes promote a state of chronic, subclinical inflammation at the inner foreskin. Members of the BASIC species directly cleave E-cadherin^8^, facilitating virion penetration^46^ and potentially contributing to local inflammation^45^. Anaerobes may also stimulate pattern-recognition pathways in keratinocytes and resident immune cells^47,48^, consistent with increased local concentrations of IL-1α, IL-1β, IL-8, and other inflammatory mediators. This inflammatory environment may in turn promote vascular activation (VEGF and MMP9) and enrichment of CCR5+/CD4+ T cells and myeloid cells within the foreskin^10,49–52^. Once there, dendritic cells and macrophages would be expected to be activated, including CD11c+ dendritic cells, enhancing their ability to capture virus at the mucosal surface and to trans-infect T cells, as described in other anogenital tissues^18,19,23,53,54^. These findings provide a mechanistic framework for how the penile microbiome influences HIV acquisition and suggest a mechanism by which MMC is protective. Targeting genital anaerobes^7,55^ remains a promising therapeutic opportunity to reduce transmission of HIV and other sexually transmitted viruses.

## Study Limitations

By combining machine-learning-based image analysis with manual refinement, we quantified HIV target cells across full-section immunofluorescence images from a relatively large number of participants and mapped their spatial localization. This approach enabled detection of CD11c+ dendritic cells, which can be lost during enzymatic tissue processing for flow cytometry^19^; however, conventional four-colour immunofluorescence did not permit full resolution of macrophage and dendritic-cell subsets, or other potential HIV target cells such as mast cells^56^. Antimicrobial treatment assignment in the parent randomized trial was also a potential confounder. Although associations between BASIC species and HIV target-cell densities remained after adjustment for treatment assignment, residual confounding by unmeasured factors – including undetected STIs, sexual behaviours, or penile taxa not captured within the BASIC category – cannot be excluded. Finally, because foreskin tissue analysis was cross-sectional and based on two-dimensional sections, we could not directly establish causality, fully resolve three-dimensional cell positioning, or measure ex vivo infection of foreskin myeloid cells. Complementary longitudinal, multiplex-imaging, and functional studies will be needed to define the mechanisms underlying these associations and assess generalizability to other populations.

## Acknowledgements

This work was supported by the Canadian Institutes of Health Research (TMI-138656 and PJT-1806; RK), the NIH National Institute for Allergy and Infectious Diseases (R01AI123002-01A1; CML, R01AI128779, AART), the NIH National Institute for Diabetes and Digestive and Kidney Diseases (R01DK131936-01; CML). RMG was supported by the NIH Fogarty HIV Research Training Program (4D43TW009578-04); JLP was supported by a Canada Research Chair; LBB was supported by the Canadian Institutes of Health Research Canada Graduate Scholarship Program. This work was conducted in part through support of The Canadian Foundation for Innovation John R. Evans Leaders Fund and the Ministry of Colleges and Universities Ontario Research Fund – Small Infrastructure Fund. The funders had no role in the design of the present analysis, data collection, data analysis, interpretation, manuscript preparation, or decision to submit the manuscript for publication. This work was made possible by the use of equipment in the Molecular Imaging Facility at Western University.

## Data Availability

Datasets are available from the corresponding author pending research ethics board approval.

## Competing Interests

The authors declare no competing interests.

## Conflict-of-Interest Statement

The authors declare no conflict of interest

## Supplemental Tables & Figures

**Supplemental Figure 1.**
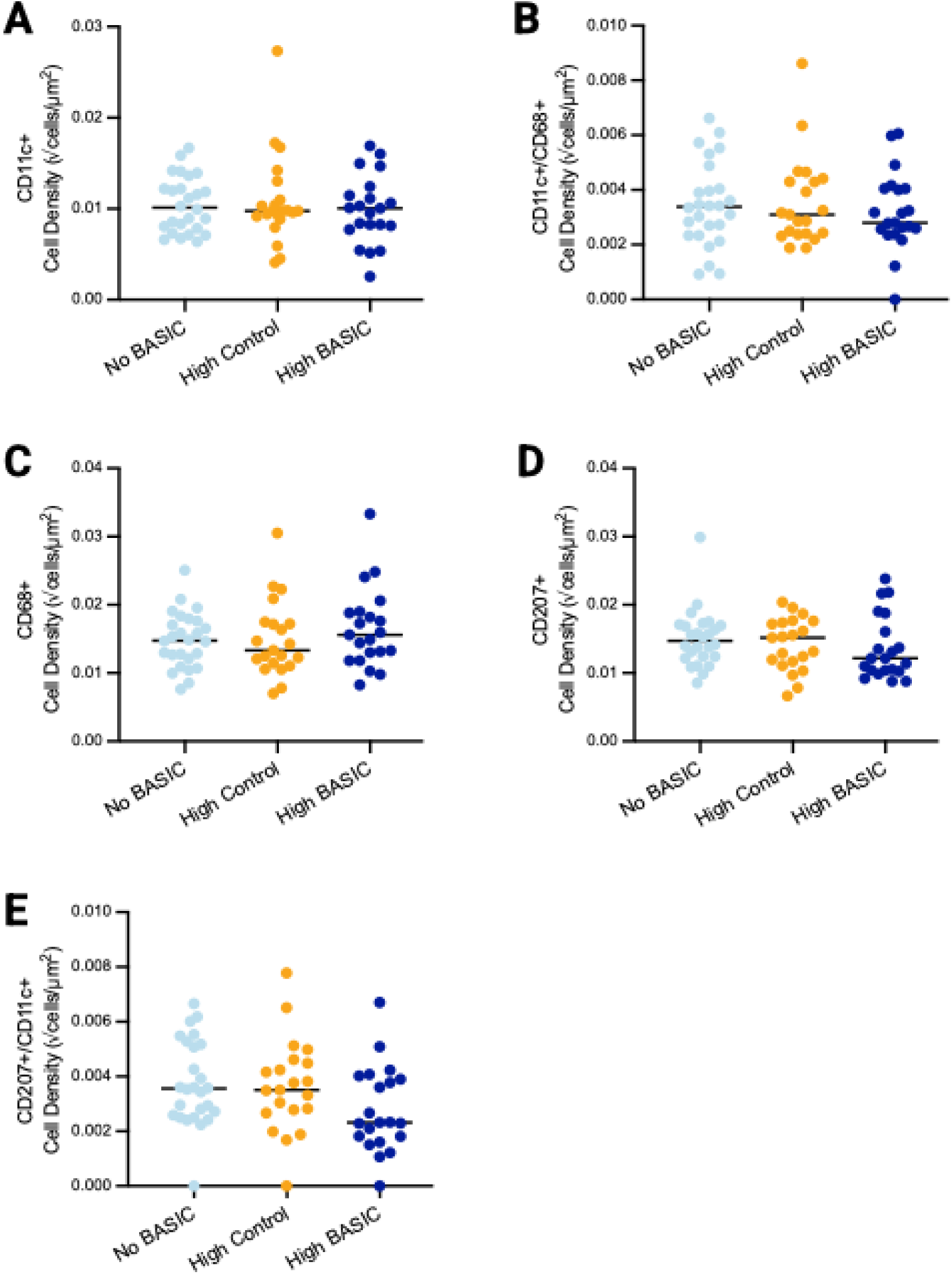
Immune cell densities in the inner foreskin epidermis. CD11c+ (**A**), CD68+ (**B**), CD207+ (**C**), CD11c+CD68+ (**D**) co-expressing, and CD207+CD11c+ (**E**) co-expressing cells compared in the epidermis across High BASIC (n=21), High Control (n=21), and No BASIC (n=26) groups. Horizontal plot lines represent the median cell density (cells/µm²) for each group. Statistical significance was assessed using the Kruskal-Wallis test followed by Dunn’s post-hoc test with Bonferroni correction, α = 0.05. No significant differences were observed.

**Supplemental Table 1.** Antibodies used for immunofluorescence and histology.

| Target | 1°/<br>2° | Clone | Supplier | Host<br>Species | Dilution | Fluorophore |
| --- | --- | --- | --- | --- | --- | --- |
| CD3 | 1° | SP7 | Abcam | Rabbit | Neat | None |
| CD4 | 1° | Polyclonal | R&D | Goat | 1:10 | None |
| CCR5 | 1° | * | * | Mouse | 1:10 | None |
| CD207 | 1° | Polyclonal | R&D | Goat | 1:20 | None |
| CD11c | 1° | EP1347Y | Abcam | Rabbit | 1:250 | None |
| CD68 | 1° | C68/684 | Abcam | Mouse | 1:500 | None |
| Mouse IgG<sup>†</sup> | 2° | Polyclonal | Fisher | Donkey | 1:400 | Alexa Fluor |
| Rabbit IgG<sup>†</sup> | 2° | Polyclonal | Fisher | Donkey | 1:400 | Alexa Fluor |
| Goat IgG<sup>†</sup> | 2° | Polyclonal | Fisher | Donkey | 1:400 | Alexa Fluor |

<sup>†</sup> Binds both heavy and light IgG chains (H + L)
\* Monoclonal hybridoma generously provided by Dr. Matthias Mack (University of Regensburg, Germany)

**Supplemental Table 2.** Excitation & Emission Filters for Immunofluorescence Microscopy.

| Target | Leica Filter | Absorbance $\lambda$ | Excitation<br>Filter<sup>a</sup> | Emission $\lambda$ | Emission<br>Filter<sup>a</sup> |
| --- | --- | --- | --- | --- | --- |
| DAPI | CFP | 358 | 436/20 | 461 | 480/40 |
| Alexa Fluor 488 | GFP | 494 | 425/60 | 517 | 480 LP |
| Alexa Fluor 546 | DSR | 556 | 545/30 | 573 | 620/60 |
| Alexa Fluor 647 | Y5 | 650 | 620/60 | 665 | 700/75 |

LP long pass filter
$\lambda$ wavelength, in nanometers
<sup>a</sup>Peak wavelength of light that passes through the filter and filter bandwidth, in nanometers

## Notes

### Competing Interest Statement

The authors have declared no competing interest.

